# GMSE: an R package for generalised management strategy evaluation

**DOI:** 10.1101/221432

**Authors:** A. Bradley Duthie, Jeremy J. Cusack, Isabel L. Jones, Jeroen Minderman, Erlend B. Nilsen, Rocío A. Pozo, O. Sarobidy Rakotonarivo, Bram Van Moorter, Nils Bunnefeld

## Abstract

1. Management strategy evaluation (MSE) is a powerful tool for simulating all key aspects of natural resource management under conditions of uncertainty.
2. We present the R package GMSE, which applies genetic algorithms to provide a generalised tool for simulating adaptive decision-making management scenarios between stakeholders with competing objectives under complex social-ecological interactions and uncertainty.
3. GMSE models can be agent-based and spatially explicit, incorporating a high degree of realism through mechanistic modelling of links and feedbacks among stakeholders and with the ecosystem; additionally, user-defined sub-models can also be incorporated as functions into the broader GMSE framework.
4. We show how GMSE simulates a social-ecological system using the example of an adaptively managed waterfowl population on an agricultural landscape; simulated waterfowl exploit agricultural land, causing conflict between conservation interests and the interest of food producers maximising their crop yield.
5. The R package GMSE is open source under GNU Public License; source code and documents are freely available on GitHub.

## Introduction

Many global natural resources, including the biodiversity on which critical ecosystem services depend, are in a state of severe decline (Dirzo et al., 2014; Hautier et al., 2015; Ceballos et al., 2017; O’Connell, 2017). Conservation of biodiversity can be complicated by the immediate need to use natural resources and land area for human livelihood, causing real or perceived conflicts between biodiversity conservation and agricultural production. This creates a challenging situation for the management of many natural resources (Redpath et al., 2015). Given increasing human population size (Crist et al., 2017), the number and intensity of such conflicts are likely to increase into the twenty first century. Effective management tools are therefore needed for the long-term sustainable use of natural resources under the rising demand for food production (Fischer et al., 2017).

To effectively manage natural resources, an adaptive approach allows managers to iteratively update their models and respond flexibly to changing conditions (Keith et al., 2011). This approach is especially effective when considering multiple aspects of the social-ecological system being managed, including the dynamics of resources, monitoring, and the decision-making processes of stakeholders (Bunnefeld et al., 2011; Bunnefeld and Keane, 2014). Management strategy evaluation (MSE) is a modelling framework, first developed in fisheries (e.g., Sainsbury et al., 2000; Polacheck et al., 1999; Smith et al., 1999; Moore et al., 2013), for simulating all of these aspects of resource management in a way that uniquely considers the uncertainties inherent to every stage of the management process (Bunnefeld et al., 2011; Punt et al., 2016). Nevertheless, MSE models developed hitherto have been limited in their ability to model human decision-making (Fulton et al., 2011; Dichmont and Fulton, 2017); manager decisions are typically based on fixed rules, and user behaviour likewise remains fixed over time instead of dynamically responding to changing resource availability and management decisions (Schlüter et al., 2012; Melbourne-Thomas et al., 2017). Here we introduce generalised management strategy evaluation (GMSE), which incorporates a game-theoretic perspective to model the goal-oriented, dynamic decision-making processes of stakeholders.

The R package ‘GMSE’ is a flexible modelling tool to simulate key aspects of natural resource management. GMSE offers a range of parameters to simulate resource dynamics (primarily, but not necessarily biological populations) and management policy options, and includes genetic algorithms to dynamically model stakeholder (manager and user) decision-making. Genetic algorithms find adaptive solutions to any simulated conditions given stakeholder-specific goals (see SI1), allowing GMSE to model scenarios of conservation conflict.

GMSE allows researchers to address adaptive management questions *in silico* through simulation. Simulations can be parameterised with initial conditions derived from empirical populations of conservation interest to predict key social-ecological outcomes (e.g., resource extinction, agricultural yield) given uncertainty. The sensitivity of these outcomes to different management options (e.g., population target, policies available, observation methods, budget constraints, etc.) can thereby inform management decisions, even given competing management objectives caused by conservation conflict (e.g., Strand et al., 2012; Redpath et al., 2013; Sundt-Hansen et al., 2015; Pozo et al., 2017; Fox and Madsen, 2017). Additionally, GMSE can be used to explore general questions concerning management theory such as the following: How is population persistence affected by management frequency or observation intensity? How does variation in user actions affect the distribution of resources or landscape properties? How do asymmetries in stakeholder influence (i.e., budgets) affect resource dynamics?

### GMSE model structure

GMSE builds off of the MSE framework (Fig. 1). The function gmse runs simulations using four predefined individual-based submodels, which can be parameterised to fit various case studies; more tailored submodels can also be defined using the `gmse_apply` function (see SI2). By default, GMSE models (1) a population of discrete resources (e.g., a managed species) with individual traits (e.g., location, age) on a spatially-explicit landscape and simulates resource birth, individual movement between landscape cells, interaction with the landscape, and death; the discrete nature of resources causes demographic stochasticity, and therefore uncertainty. This sub-model is unique in not relying on other sub-models because ecological dynamics can be simulated in the absence of observation and management. (2) Observation is modelled in one of three ways: resource counting on a subset of landscape cells (e.g., Nuno et al., 2013), marking and recapturing a fixed number of resources, or counting resources across the whole landscape one set of landscape cells at a time (during which resources might move). Sampling error from all observation types generates a range of uncertainties that depend on monitoring effort. (3) Managers analyse data collected from observations to estimate resource abundance, then compare this estimate with their pre-defined target abundance. Policy is developed by calling the genetic algorithm (see below), which works within a manager’s constraints to find costs for user actions on the resource (e.g., culling, scaring, etc.) that minimise deviation from the target abundance, as informed by the predicted consequences of each action on resource abundance and user action histories. After a suitable policy is found, (4) users perform actions that affect resources or landscape cells (e.g., culling, which causes immediate resource death). Users respond to policy individually, each calling the genetic algorithm to find actions that maximise their own utilities (e.g., maximise resource use or landscape yield) within their imposed constraints. Once each user has found an adaptive strategy, user actions affect resources and landscape cells, feeding back into the resource sub-model.

**Figure 1:**
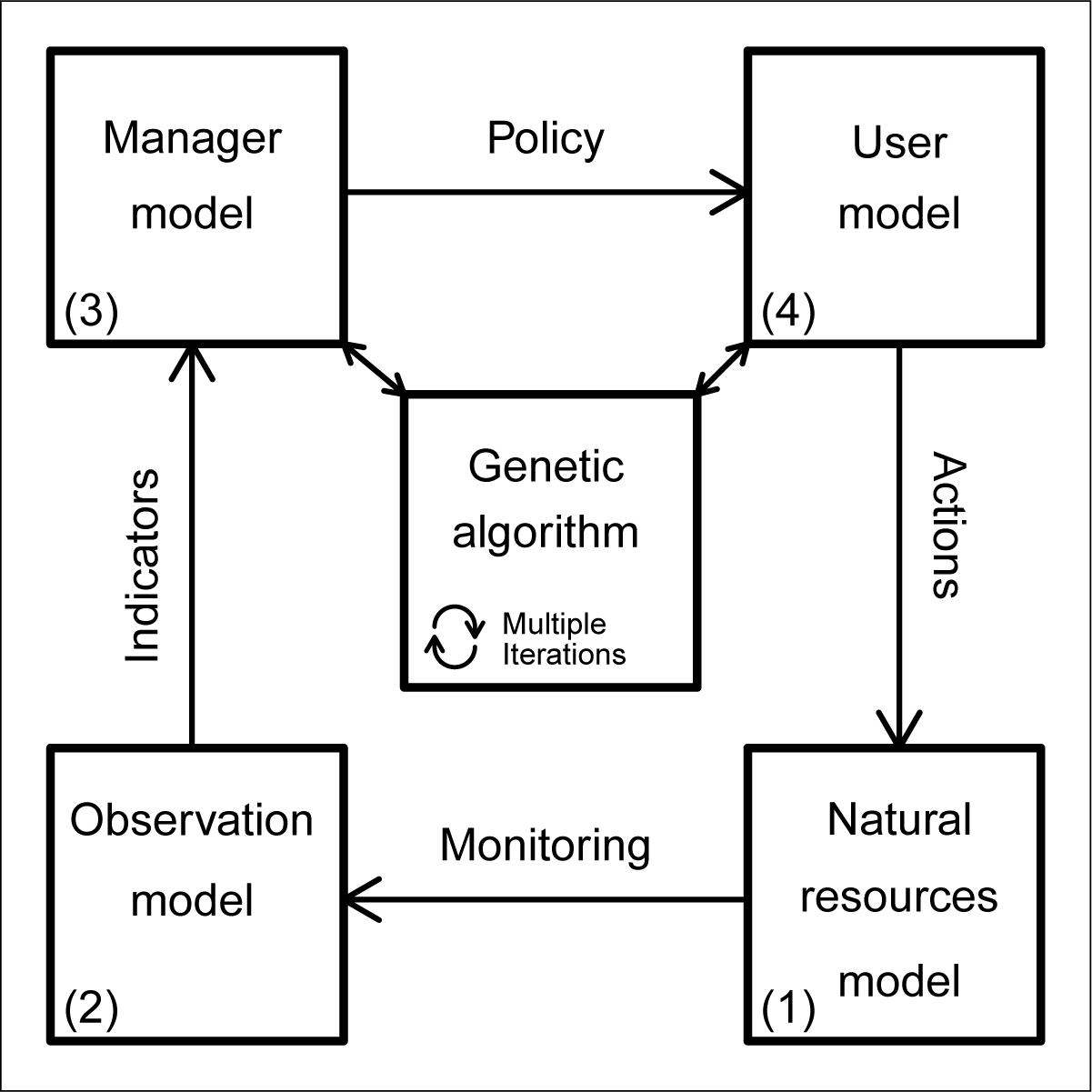
Description of one time step of the generalised management strategy evaluation framework, which is comprised of four separate sub-models.

### Genetic Algorithm

Consistent with the MSE approach (Bunnefeld et al., 2011), GMSE does not attempt to find optimal strategies or solutions for agents (stakeholders). Instead, a genetic algorithm is used to heuristically find strategies that reflect the individual objectives of each stakeholder in each time step (see SI1 for details). Critically, all stakeholders involved in resource conservation are constrained in their decision-making; managing and using resources takes effort (e.g., time or money), and effort expended in developing or enforcing one policy (for managers) or performing one action (for users) will be effort not expendable elsewhere (Milner-Gulland, 2011; Müller-Hansen et al., 2017; Schlüter et al., 2017). In finding strategies, GMSE models this trade-off by setting a fixed budget for managers and users. Allocations from a manager’s budget can be used to increase the cost it takes a user to perform an action (i.e., ‘policy’), and allocations from a user’s budget can be used to perform the action at the cost set by the manager. Hence, stakeholders can have incomplete control over resource use and express competing management objectives, potentially resulting in conflict.

A single call of the genetic algorithm simulates the process of thinking and decision-making for one manager or user. In each new call of the genetic algorithm, a unique population of temporary manager or user strategies is initialised. In each iteration of the genetic algorithm, these strategies crossover and mutate; when this results in strategies that are over-budget, expenditures are iteratively decreased at random until budget constraints are satisfied. A fitness function then evaluates each strategy in the population, and a tournament is used to select the next iteration of strategies (Hamblin, 2013). The genetic algorithm terminates when a minimum number of iterations has passed and the increase in the fitness of the fittest strategy between the current and previous iteration is below some threshold. The highest fitness strategy in the population then becomes the stakeholder’s new strategy.

### An example of resource management

Here we illustrate the usefulness of GMSE by considering the case study of a protected population of waterfowl that exploits agricultural land causing a conservation conflict with farmers (e.g., Fox and Madsen, 2017; Mason et al., 2017; Tulloch et al., 2017). Managers attempt to keep the abundance of waterfowl at a target level, while farmers attempt to minimise the damage inflicted on their crops (e.g., Madsen et al., 2017). Using GMSE, we can simulate waterfowl population dynamics, along with the continued monitoring and policy set by managers, and the actions that farmers take to protect their crop yields given the constraints of policy. We consider a population of waterfowl with an initial abundance and manager target abundance of 200, but whose carrying capacity is 2000. Waterfowl consume and destroy all crop yield upon arrival to a landscape cell. In each time step, waterfowl are observed on a random subset of cells, then managers extrapolate from density per cell to estimate total population size. Managers then use these estimates to set costs of culling and scaring waterfowl for five farmers (scaring is non-lethal, causing waterfowl to move to a random landscape cell). Farmers attempt to reduce the negative impact of waterfowl on the cropland that they own, working within the constraints of culling and scaring costs and their budget for performing these actions.

~~~
sim <- **gmse**(land_ownership = TRUE, stakeholders = 5, observe_type = 0,
            res_death_K = 2000, manage_target = 1000, RESOURCE_ini = 200,
            user_budget = 1000, manager_budget = 1000, res_consume = 1,
            scaring = TRUE, plotting = FALSE);
## [1] “Initialising simulations … ”
## [1] “Generation 27 of 100”
## [1] “Generation 56 of 100”
## [1] “Generation 85 of 100”
~~~

Parameters in `gmse` not listed are set to default values. By plotting the output with `plot_gmse_results`, simulation results can be interpreted visually (manager and user decisions can also be interpreted using the `plot_gmse_effort` function, see SI3 and SI4 for an expanded example).

~~~
**plot_gmse_results**(sim_results = sim);
~~~

Fig. 2 shows the landscape broken down by resource position and farmer land ownership in the upper left and right hand panels, respectively. The waterfowl population fluctuates around the manager’s target size of 1000, but the manager’s estimate of population size deviates from its actual size due to observation uncertainty (compare black and blue lines in the middle left panel). Because the waterfowl have a direct negative effect on landscape yield, total landscape yield (orange line of the middle left panel), along with the yield of individual farmers (right middle panel), is low when waterfowl abundance is high, and vice versa.

**Figure 2:**
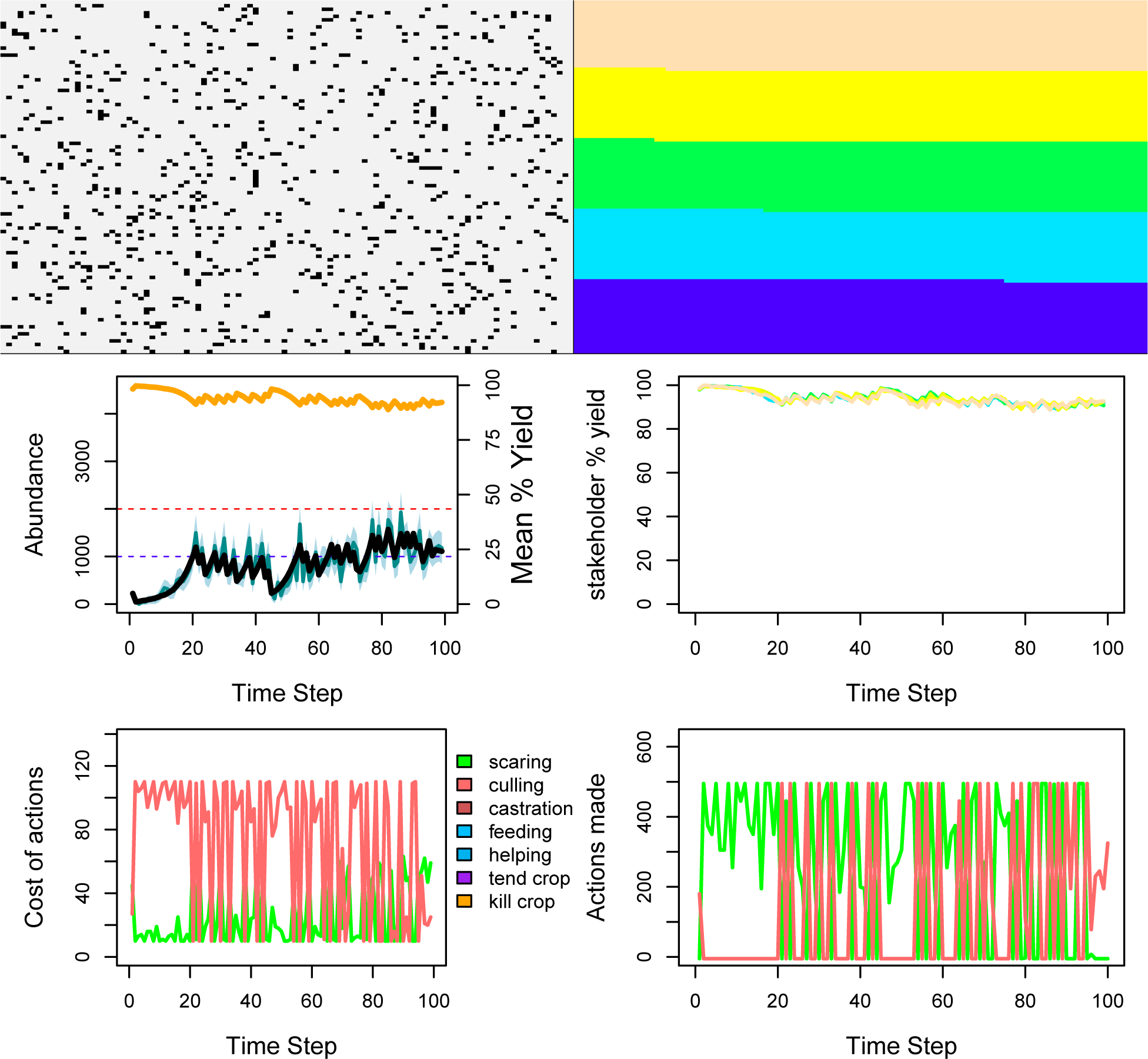
Results of an example simulation illustrating the management of a protected resource that exploits the land of five farmers. The upper left panel shows locations of resources (black dots) on the landscape in the final time step of the simulation (multiple resources can occur on the same landscape cell). The upper right panel shows the same landscape broken down into five differently coloured regions, which correspond to areas of land owned by each of the five farmers. The middle left panel shows the actual abundance of resources (black solid line; i.e., ‘natural resources’ or ‘operating’ model), and the abundance of resources as estimated by the manager (blue solid line; i.e., ‘observation’ or ‘assessment’ model; shading indicates 95 percent confidence intervals), over time. The horizontal dotted red and blue lines show the landscape-level resource carrying capacity enacted on adult mortality and the manager’s target for resource abundance, respectively. The orange line shows the total percent yield of landscape cells. The middle right panel shows total percent yield of landscape cells for each individual farmer, differentiated by colour, where line colours correspond to areas of the landscape in the upper right panel. The lower left panel shows the cost of farmers performing actions over time, as set by the manager; the upper limit on cost of actions reflects the manager’s limited budget for setting policy. The lower right panel shows the total number of actions attempted to be performed by all farmers over time (some actions might be unsuccessful if resources are not on a farmer’s land to cull or scare, so, e.g., culling actions might be larger than resources actually culled).

Only the estimates of population size from the observation model are available to the manager, so policy change at any time step is driven primarily by the deviation of the currently estimated population size from the manager’s target and the actions of farmers in the previous time step. Hence, when the population size is estimated to be below (above) the manager’s target, the manager increases (decreases) the cost of culling and decreases (increases) the cost of scaring. Because the manager does not know in advance how farmers will react to policy change, they assume a proportional response in total actions with respect to a change in cost (e.g., doubling the cost of culling will decrease stakeholder culling by 1/2). Farmers responding to policy are interested only in minimising waterfowl’s exploitation of their crops, so they will either cull or scare to remove the waterfowl from their land, depending on which option is more effective (i.e., cheaper). This is reflected in the bottom left versus right panels of Fig. 2; when managers decrease culling costs relative to scaring, farmers respond with more total culling, and vice versa. Farmer decisions then affect waterfowl distribution and abundance, impacting future crop yield and policy.

### Graphical User Interface (GUI)

The function gmse_gui opens GMSE in a browser and allows simulations to be run for most `gmse` parameter options using the package ‘shiny’ (Chang et al., 2017). Figures from `plot_gmse_results` and `plot_gmse_effort`, and tables from `gmse_summary` are provided as GUI output.

### Custom defined sub-models

The function `gmse_apply` allows custom resource, observation, manager, or user sub-models to be integrated into the GMSE framework (see SI2, SI4, and SI5). Any type of sub-model (e.g., numerical, individual-based) is permitted by defining a function with appropriately specified inputs and outputs; where custom functions are not provided, `gmse_apply` runs default GMSE sub-models used in `gmse`. Any parameter options available in `gmse` or in custom functions can be passed directly to `gmse_apply`, thereby allowing for high flexibility in model specification. For example, a simple logistic growth function can be integrated as a resource sub-model to replace the default `resource` function.

~~~
logistic_res_mod <-function(X_0, K = 2000, gr = 1){
    X_1 <-X_0 + gr * X_0 * (1 - X_0/K);
    **return**(X_1);
}
sim <- **gmse_apply**(res_mod = logistic_res_mod, X_0 = 200, gr = 0.3, stakeholders = 5);
~~~

The `gmse_apply` function simulates a single GMSE time step, and therefore must be looped for simulations over multiple time steps. Within loops, GMSE arguments can be redefined to simulate changing conditions (e.g., change in policy availability or stakeholder budgets, see SI2 and SI4), thereby allowing many management scenarios to be simulated *in silico*.

### Availability

The GMSE package can be downloaded from CRAN (https://cran.r-project.org/package=GMSE) or GitHub (https://confoobio.github.io/gmse/). GMSE is open source under GNU Public License.

## Conclusions

Here we have introduced the R package GMSE v0.4.0.3, new software for modelling social-ecological dynamics under scenarios of potential conflict. GMSE provides a powerful tool for individual-based modelling simulations while also allowing for extensive model customisation. GMSE vignettes provide additional examples for getting started with `gmse` or `gmse_gui` for simulations using default resource, observation, manager, and user sub-models (SI3 and SI6), and for using `gmse_apply` for advanced model customisation (SI2, SI4, and SI7) and integration with existing packages (e.g., Fisheries Library in R, see SI5). Future versions of GMSE will include additional features and improve upon the realism of social and ecological modelling components, while also maintaining a high degree of flexibility and modulatarity for model customisation.

## Acknowledgements

This project was funded by the European Research Council under the European Union’s H2020/ERC grant agreement no. 679651 (ConFooBio) to NB. ABD is funded by a Leverhulme Trust Early Career Fellowship. We thank two anonymous reviewers for helpful comments.

